# Cave Adaptation Favors Aging Resilience in the Mexican Tetra

**DOI:** 10.1101/2024.09.26.615235

**Authors:** Ansa E. Cobham, Alexander Kenzior, Pedro Morales-Sosa, Jose Emmanuel Javier, Selene Swanson, Christopher Wood, Nicolas Rohner

**Affiliations:** Stowers Institute for Medical Research, Kansas City, MO 64110; Department of Cell Biology & Physiology, University of Kansas Medical Center, Kansas City, Kansas, USA

## Abstract

All animals age, but the rate at which some species age varies widely. What environmental pressures and what molecular factors underlie the remarkable diversity in aging and lifespan across species remains largely enigmatic. The Mexican tetra, *Astyanax mexicanus*, serves as an intriguing new model to study how adaptations to different environments can change aging. This species exists as two morphotypes: the river-dwelling surface fish, which inhabits environments rich in food and light, and the blind cave-adapted cavefish, that thrive in dark, nutrient-limited but predator free environments. Nutrient limitation and lack of predation are known to impact lifespan, however, how adaption to such environments changes the aging trajectory in this species remains unknown. Here, we compared aging markers between surface and cavefish populations, focusing on morphological and behavioral changes, as well as molecular signatures, and found that aging markers are pronounced and evident in surface fish, whereas these signatures are less distinct in aged cavefish. Using zebrafish, we explored the contribution of the cavefish insulin receptor mutation to longevity. Although the insulin receptor mutation is sufficient to increase lifespan in fish, our findings suggest its impact is limited. Instead, our data indicate that metabolic shifts, particularly those related to mitochondrial function, may be key contributors to the extended longevity observed in cavefish.

## Introduction

Aging is a complex biological process characterized by a gradual decline in physiological functions, ultimately leading to death. This inevitable aspect of life has captivated humanity throughout history, driving many to seek ways to halt or reverse this process. Although aging is a ubiquitous phenomenon among species, suggesting a highly conserved biological underpinning, there exists remarkable diversity in lifespan and aging patterns across the animal kingdom. Generally, larger animals tend to live longer than smaller ones, though numerous exceptions exist. For instance, rats typically live about three years, whereas squirrels and naked mole-rats can thrive for up to twenty-five years (Gorbunova et al., 2008). Outside of mammals, there exists an even wider variety, for example, some sea urchin species can live for centuries, while certain fish live only long enough to reproduce before dying (Ebert, 1982; Healey, 1991).

The reason why evolution favors variations in longevity is largely unknown. Predator-free environments, such as caves, often harbor species with longer lifespans than their surface-dwelling counterparts. However, pinpointing species-specific aging signatures and linking them to selective pressures is challenging, due to the lack of a consensus on common aging indicators across species. Signs of aging include loss of reproductive capacity, morphological decay, cellular senescence, telomere attrition, and others, each contributing uniquely to the aging narrative across species (Lopez-Otin et al., 2013, 2023). Furthermore, aging tends to increase biological vulnerability to stressors, leading to a decreased capacity to maintain physiologic homeostasis and thereby increasing the risk of mortality (Fried et al., 2004; Vaupel et al., 1979).

Some species appear to mitigate some of the detrimental effects of aging. Evidence from long-lived vertebrate species indicates a combination of longevity-favoring traits, such as increased antioxidant and repair systems, enhanced regenerative capacity, metabolic slowdown, and maintenance of lower body temperatures to minimize oxidative stress (Shilovsky et al., 2022). Fasting is among the most researched anti-aging interventions; however, the mechanisms by which caloric restriction extends lifespan are not fully understood. The Insulin and Insulin-Like Growth-Factor Signaling pathway is a prime candidate for lifespan extension (Kenyon, 2005). Multiple evidence shows that manipulating the IIS pathway to reduce insulin activity (insulin resistance) consistently extends lifespan in worms, flies, and mammals (Gresl et al., 2003; Tatar et al., 2003). A single mutation reducing insulin signaling is sufficient to double lifespan in *C. elegans* and *D. melanogaster* (Kenyon, 2005; Kenyon et al., 1993). However, this effect is less pronounced in mice and other mammals. Long-lived mouse models with markedly reduced insulin signaling under dietary restriction often display increased insulin resistance and increased glucose tolerance (Davidson et al., 2002; Kalant et al., 1988; McCarter et al., 2007). Additionally, reduced insulin sensitivity is a risk factor for hyperglycemia and diabetes (Barzilai et al., 2012), making it counterintuitive that insulin resistance would extend lifespan. These paradoxes on the link between insulin resistance and aging raise questions about whether the IIS pathway alone is sufficient as an evolutionarily conserved modulator of aging. Thus, identifying alternative vertebrate models holds promise for shedding light on insulin signaling and other specific regulators of aging.

In this study, we present the Mexican Tetra, *Astyanax mexicanus*, as a unique study system for exploring the intersection of environmental factors and metabolic changes during aging. This species exhibits two major morphs—surface and cave-dwelling—that significantly differ in selective pressures affecting survival, metabolism, and insulin signaling (Cobham & Rohner, 2024; Medley et al., 2022; Olsen et al., 2021; Riddle et al., 2021). The surface fish inhabit environments rich in food but fraught with predators, while the cave-adapted forms reside in almost predator-free zones with limited or seasonally restricted food supplies, conditions that theoretically favor longer lifespans. Indeed, laboratory studies have shown that cavefish live longer than surface fish and exhibit a transcriptional profile similar to younger individuals of their species (Riddle 2018; Lloyd 2024). Additionally, cavefish possess a mutation in the insulin receptor that induces insulin resistance (Riddle et al., 2018), potentially linking this genetic alteration to their increased longevity.

Here, we explored morphological and behavioral signs of aging in young and aged individuals of *Astyanax mexicanus* and assessed the impact of the cavefish insulin receptor mutation on longevity using the zebrafish model. We found a slight increase in lifespan in zebrafish that were genetically engineered to carry the cavefish allele, suggesting that additional mechanisms beyond insulin signaling may be contributing to the age-defying mechanisms of cavefish. Furthermore, protein and metabolic assays indicates that metabolic shifts in mitochondrial activity may underlie age-related differences. Collectively, the molecular markers of metabolic adaptation may contribute to the aging-related resilience and enhanced health span in the cavefish.

## Results

### Morphological signs of aging are reduced in cavefish

To systematically study aging in *Astyanax mexicanus*, we compared morphometric data from 30 laboratory raised surface fish and 30 cavefish across three different age groups: young (2 years), middle-aged (6 years), and old (9 years). We measured standard length, body weight, and spine curvature (**Figure 1**).

**Figure 1.**
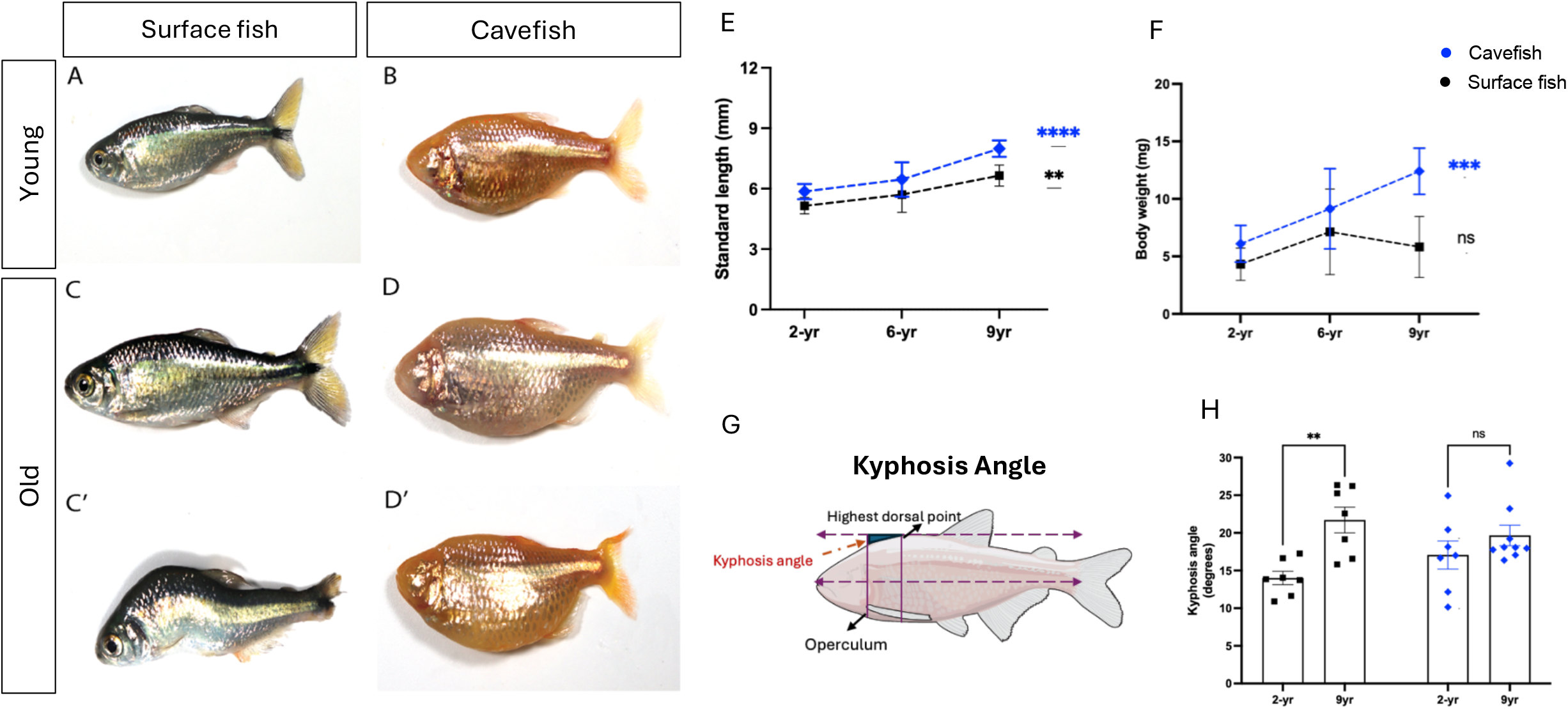
Body condition changes in aged fish. Young surface fish **(A)** and cavefish **(B)** compared with older fish **(C)** and **(D)** respectively. Visible spine deformity (kyphosis) was only observed in old surface fish **(C’)** compared to cavefish **(D’)**. Measurements of Standard length **(E)** and Body weight **(F)** indicate body wasting in old surface fish. **(G)** Graphic for quantification of spinal deformity Kyphosis angle and **(H)** Measurements of kyphosis angle across different aged cavefish and surface fish. n = 9-10 for surface and cavefish per timepoint. Data are represented as mean + SD, *P < 0.05, **P < 0.01, ***P < 0.001, ****P < 0.0001, ns = not significant.

Consistent with known patterns of continuous growth throughout their lifetimes, both morphs exhibited increased standard lengths at older ages (**Figure 1E**). Similarly, body weight also increased with age in cavefish, however in surface fish we found that the old individuals showed a 50% decrease in body weight from 6 to 9 years of age (**Figure 1F**). Such decrease in body weight has been reported to be a common sign of aging and is usually associated with body wasting (Thomas, 2007).

Given the association between kyphosis (abnormal curvature of the spine) and body wasting across several organisms including zebrafish (Gerhard et al., 2002), we quantified the kyphosis angle in *Astyanax mexicanus* (**Figure 1G**). We did not observe abnormal spine curvature in 2-or 6-year-old surface and cavefish. However, the 9-year-old surface fish showed significant degrees of kyphosis (**Figure 1H**). Intriguingly, we did not detect visible kyphosis in the old cavefish, aligning with prior observations that cavefish exhibit slower aging processes (Riddle 2018, Lloyd 2024).

### Swimming behavior is affected with age

Cavefish show a robust range of swimming patterns (Olsen et al., 2023; Patch et al., 2022; Paz et al., 2020; Tan et al., 2011). To evaluate the impact of aging on swimming performance and assess if the observed morphological changes would affect swimming behavior, we captured digital videos of young and aged cavefish and surface fish. Using animal tracking software, we measured the maximum swim velocity and total distance traveled, using frame-by-frame distances captured in each (x,y) coordinate position (**Figure 2A-E**).

**Figure 2.**
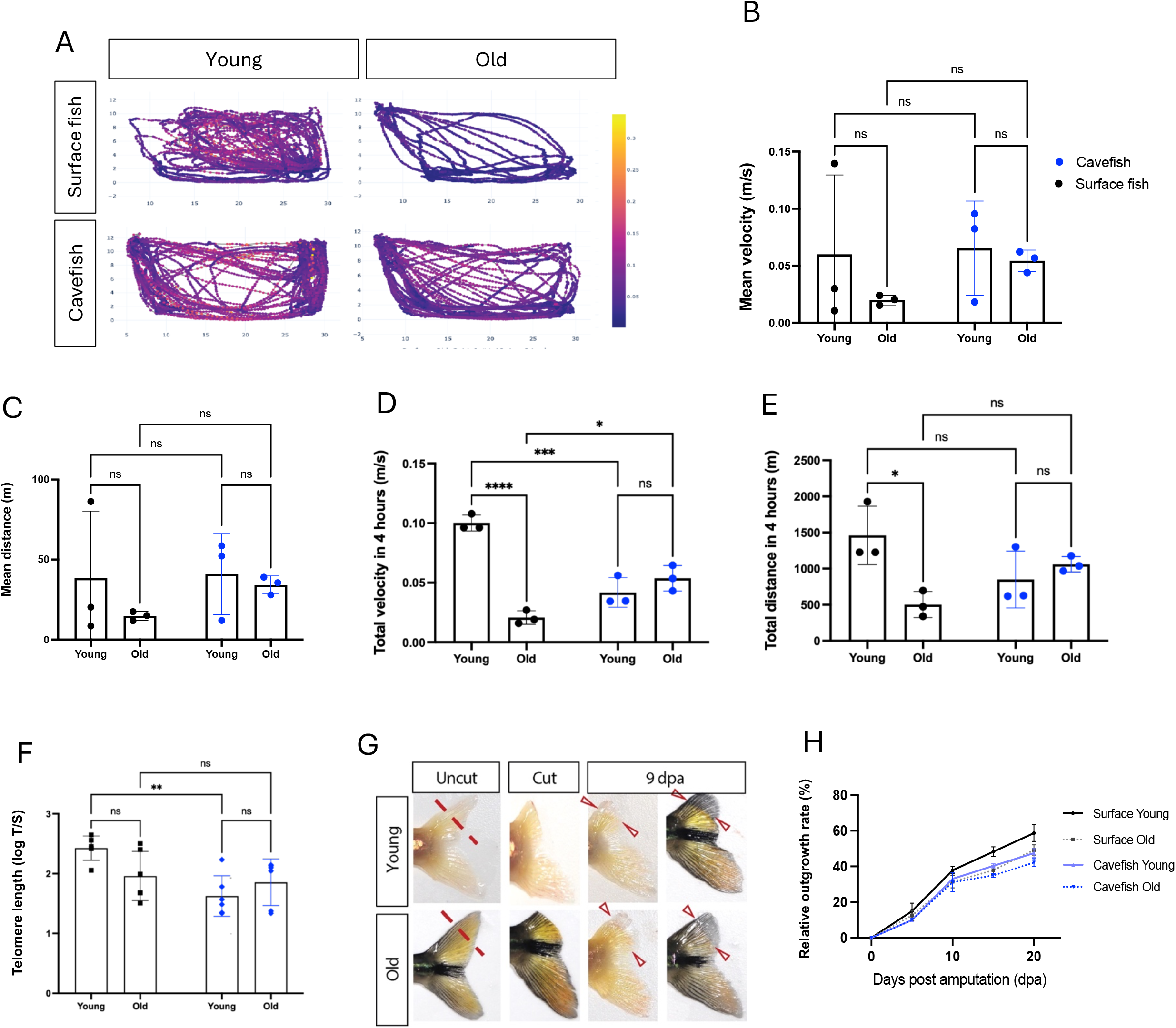
Swimming performance, telomere length, and caudal fin regeneration in aged fish. **(A)** Heat maps showing the time spent in each x and y coordinate position of young and old surface fish and cavefish over a 10-minute interval. Measurements of **(B)** mean velocity and **(C)** mean distance traveled in 10 minutes **(D)** total velocity and **(E)** total distance recorded over 4 hours. n = 3 for surface and cavefish per timepoint. **(F)** qPCR quantification of telomere length in caudal fin of cavefish and surface fish. **(G)** Caudal fin regenerates in old and young fish days post amputation (dpa). **(H)** Quantification of the relative outgrowth in the regenerated caudal fin in young and old surface and cavefish. (B-E), n = 3, (F and H), n=6-7 for surface and cavefish per timepoint. Data are represented as mean + SD, *P < 0.05, **P < 0.01, ***P < 0.001, ****P < 0.0001, ns = not significant.

Our analysis was done in two timepoints, short multiple 10 minutes intervals and total measurements, over 4-hours collected at a similar time across experiments. Within the 10 minutes interval, our analysis revealed a trend towards reduced mean velocity in aged surface fish compared to young surface fish, (0.060 versus 0.020 + 0.045m/s; p= 0.6412), however, this was not statistically significant (**Figure 2B**). Mean velocity did not change with age in the cavefish (0.065 versus 0.054 + 0.011 m/s; p= 0.9866. Similarly, mean distance traveled was not significantly reduced in both surface fish and cavefish with age (38.37 versus 14.77 + 23.60m; p=0.6600, **Figure 2C**).

Over the entire 4-hour time interval, we observed a significant difference in total velocity between surface fish and cavefish as well as in an age-dependent manner. Young surface fish had the highest speed traveled compared to the old surface fish (0.100 versus 0.021 + 0.079m/s; p=<0.0001, **Figure 2D**). This maximum speed was also significantly higher in the young cavefish (0.100 versus 0.042 + 0.058 m/s; p=0.0003, **Figure 2D**). Remarkably, however, both total velocity and total distance traveled significantly decreased with age in the surface fish (1460 versus 502.5 + 957.8m; p=0.0190), but not in the cavefish (849.7 versus 1061 + 211.3m; p=0.8246, **Figure 2D, E**).

### Telomere length and fin regeneration varies with age

To study cellular markers of aging in *Astyanax mexicanus*, we measured telomere lengths across ages in both surface and cavefish. Telomeric DNA comprises repeat sequences at the end of chromosomes, that protect animals from age-related degradation and damage (Blackburn et al., 2006; Lopez-Otin et al., 2013) and has been proposed as a good biomarker for biological age across different organisms including fish (Anchelin et al., 2011; Boonekamp et al., 2013). Remarkably, we observed that telomere length did not change significantly with age in either morph (**Figure 2F**), however, in line with previous observations (Lunghi & Bilandzija, 2024), telomere length was markedly shorter overall in cavefish than in the surface fish, suggesting that telomere length may be unrelated to delayed aging in cavefish.

Aging and telomere length are known to play critical roles in tissue repair in other teleost fish (Anchelin et al., 2011; Elmore et al., 2008; Lund et al., 2009). To establish this relationship in *Astyanax*, we performed a caudal fin regeneration assay to assess the regeneration process at the different ages. Consistent with previous results, all animals had the potential to regenerate their fins (**Figure 2G**), with rapid growth observed within the first 9-days-post-amputation (dpa) (**Figure 2G-H**). By this time, more than 30% of the fin was regenerated, with young surface fish regenerating 38% of their fin, and old surface fish achieved 31% regeneration. While the cavefish regenerated 33% and 31% of their fin in young and old fish, respectively. Although we did not observe a significant difference in regeneration between surface and cavefish, surface fish varied in their regenerative capacity with age (**Figure 2H**). Specifically, by the end of our observation at 20dpa, young surface fish achieved 60% of their fin whereas the old surface fish reached only about 49% of their original size. The young cavefish achieved 48% while the old cavefish reached 42% of their original fin size. These results underscore that while surface fish exhibit age-related decline in their fin regeneration capacity, cavefish show less of these age differences.

### Lifespan extension in Zebrafish with *insra* mutation

To study the molecular mechanisms of the observed differences between cavefish and surface fish aging, we focused on the potential role of insulin signaling in these fish. Our previous research revealed a coding mutation in the insulin receptor gene (*insra*) in cavefish, leading to insulin resistance and higher blood sugar (Riddle et al., 2018). Given that reduced insulin signaling is linked to longevity in other species, we reasoned that the mutation could underly aspects of the age protective phenotypes in cavefish.

To test this, we took advantage of the engineered zebrafish carrying the cavefish P211L *insra* allele at the endogenous locus (**Figure 3A**) (Riddle et al, 2018) to measure the effect of this mutation on lifespan in fish (**Figure 3B-G**). We observed that zebrafish carrying at least one allele of the cavefish insulin receptor mutation showed significantly extended lifespan compared to zebrafish homozygous for the wild-type (WT) allele (median lifespan = 470 days vs. 446 days; p = 0.0136) (**Figure 3B**). This finding was confirmed in another independent experiment (**Figure 3E**)

**Figure 3.**
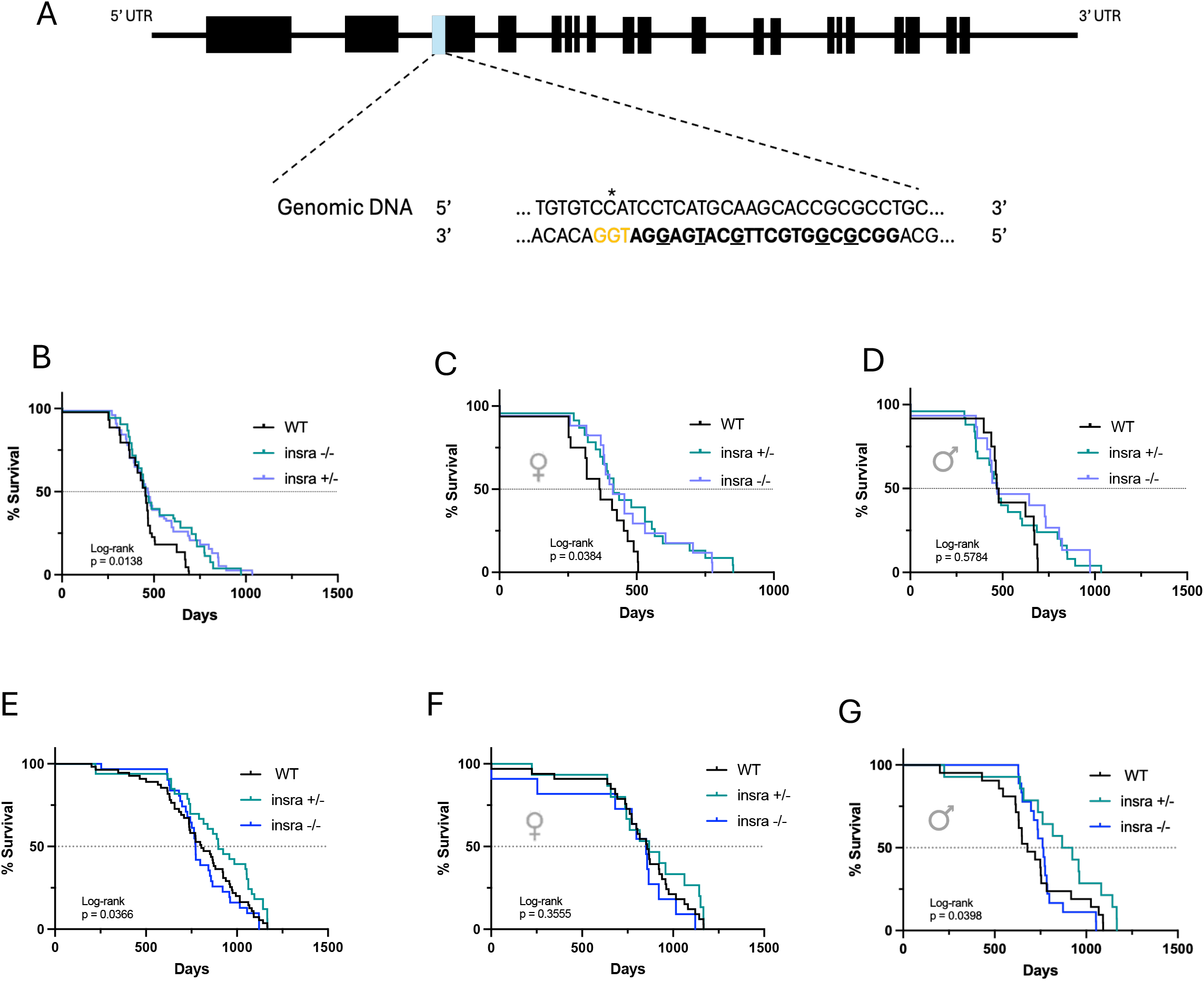
Longevity analysis of *insulin receptor a* (*insra)* mutant zebrafish. **(A)** Zebrafish P211L (*insra*) mutation using CRISPR/Cas9 strategy. Target site located in exon 3 and specific SNP exchange denoted with a star, adapted from (Riddle et al., 2018). **(B)** Survival curve of wildtype (WT), homozygote mutants (*insra* -/-) and heterozygote (*insra*+/-), in females (**C)** and males **(D**). Repeat experiment in independent cohort **(E-G)**

Insulin mutation effect on lifespan was also observed in a sex-dependent manner. In the first cohort, mutant females lived significantly longer than WT females (median lifespan = 434 days vs. 368 days; p = 0.0384) (**Figure 3C**), while this increase in lifespan was not significantly observed in either WT or mutant males (median lifespan = 474 days vs. 475 days; p = 0.5784) (**Figure 3D**). In the second experimental cohort, this sex effect was reversed. We observed that lifespan was extended in males carrying either one or two copies of the mutant allele (p=0.0398) (**Figure 3G**), while females did not have any significant increase in lifespan. Although the mechanisms that underlie aging-related insulin resistance remains unclear, we recapitulate the role of insulin in extending lifespan on zebrafish carrying the cavefish mutation on the insulin receptor. Overall, these findings highlight that while the insulin receptor mutation might play a role in increasing lifespan, this effect is unlikely to fully explain the enhanced and prolonged lifespan observed in *Astyanax* cavefish populations.

### Aging alters serum protein expression in cavefish

To identify molecular predictors associated with aging in surface fish and cavefish, we used mass spectrometry to quantitatively profile abundance of circulating proteins found in the serum of young and old cavefish and surface fish (2- and 9-years-old). Across both age groups, we reliably identified 292 unique proteins. Out of these, 109 proteins were significantly expressed in an age-specific manner, with 54 of these proteins being expressed differentially in the older groups of both surface and cavefish populations (**Figure 4A**). 13 were significantly higher in old surface fish, and 41 proteins exhibited higher expression levels in cavefish

**Figure 4.**
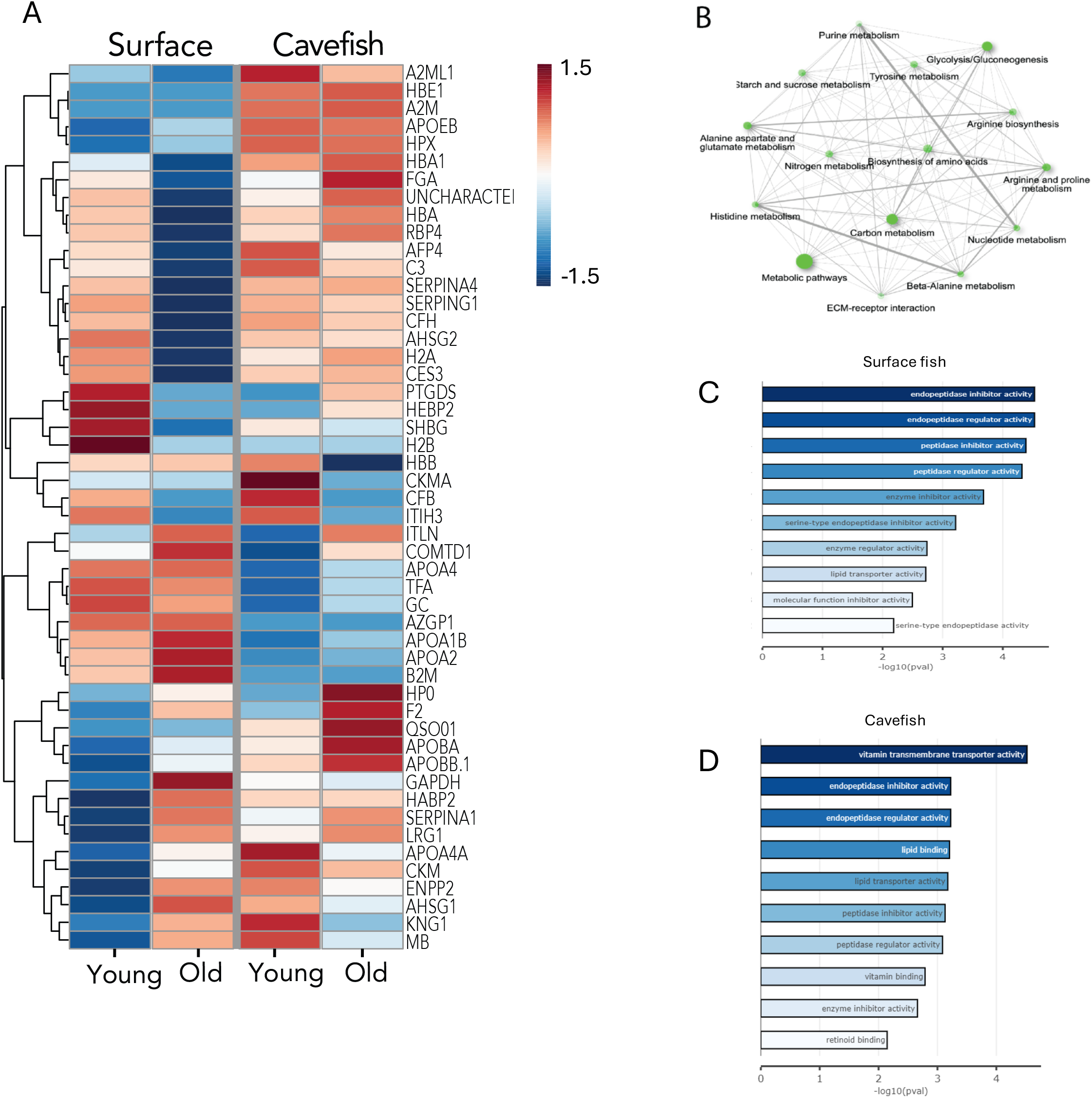
Serum proteomic analysis in cavefish and surface fish populations. (**A)** Diderential expression of proteins in young and old cavefish compared with surface fish. **(B)** Key metabolic pathways highlighted in old fish of both morphs. **(C)** Enriched pathways in old surface fish and **(D)** old cavefish. n = 3 for surface and cavefish per timepoint.

We observed that in the aged surface fish population, there was increased expression of apolipoprotein A subclasses (APOA1b, APOA2 and APOA4A), which play crucial roles in lipid transportation and lipoprotein metabolism. Additionally, peptides involved in iron regulation and oxygen transport, such as Transferrin-a (TFA) and myoglobin (MB), showed increased expression levels in aged surface fish. Similarly, levels of Catechol-O-Methyltransferase Domain (COMTD) and the glycolytic enzyme GAPDH were also higher in older fish compared to the young fish. The increase in abundance of these protein levels has been associated with age-related elevation in oxidative stress levels (Lowe et al., 2000; Nicholls et al., 2012; Tian et al., 2022).

In the cavefish population, proteins associated with lipid metabolism, peptidase regulation and antioxidant function showed increased expression levels between young and old fish. These include SERPINE1, APOB, APOE, HPX, LRG1 and IGLC1. Interestingly, LRG1 has been implicated in slowing down age-related tissue decline through extracellular matrix remodeling and neovascularization of tissues (Camilli et al., 2022; Park et al., 2023). It also acts as an adipokine that mediates systemic anti-inflammatory effects and promotes insulin sensitivity in humans and mice (Choi et al., 2022).

To establish the biological relevance of these changes, we queried gene ontology (GO) and Kyoto Encyclopedia of Genes and Genomes (KEGG) databases and showed enrichment of proteins in metabolic pathways (**Figure 4B**). Functional annotation showed that specific pathways enriched in the old surface fish include lipid transport and protease inhibition activities (**Figure 4C**). On the other hand, pathways enriched in the aged cavefish are involved in vitamin binding activities and endopeptidase regulation (**Figure 4D**). Together, this proteomic analysis suggests that while surface fish show markers of age-related tissue decline, cavefish show protective markers from age-related tissue decline.

### Aging leads to metabolic reprogramming in cavefish

Given that our proteomic analysis revealed age-related differences in metabolic regulation between cave and surface fish, we proceeded to evaluate the global metabolic activities within the liver, which serves as the key metabolic hub connecting energy to various tissues. Extensive evidence shows that the progressive reduction in energy production with age is primarily due to a decline in mitochondrial function (Bratic & Trifunovic, 2010; Harman, 1972), and the disruption of homeostasis in major biological fuels, including fatty acids, carbohydrates and amino acids (Barzilai et al., 2012; Voet & Voet, 2010).

Using a variation of the flow cytometry-based method SCENITH™ (Arguello et al., 2020) (**Figure 5A**) which utilizes O-propargyl-puromycin (OPP), we quantified protein translation at a single-cell level from liver tissue. This method indirectly measures relative ATP production by measuring overall protein synthesis through OPP integration. (Arguello et al., 2020) demonstrated that treating cells with inhibitors of either glucose metabolism or mitochondrial respiration directly impacts protein translation, which can be measured through the mean fluorescence intensity (MFI) of OPP integrated into overall intracellular protein. The amount of OPP is measured through click chemistry with a Picolyl azide conjugated to Alexa Fluor 647. This method allowed us to assess 1) glycolysis vs mitochondrial respiration and 2) glucose dependence vs fatty acid and amino acid oxidation (FAAO) to generate ATP in liver cells from young and aged animals.

**Figure 5.**
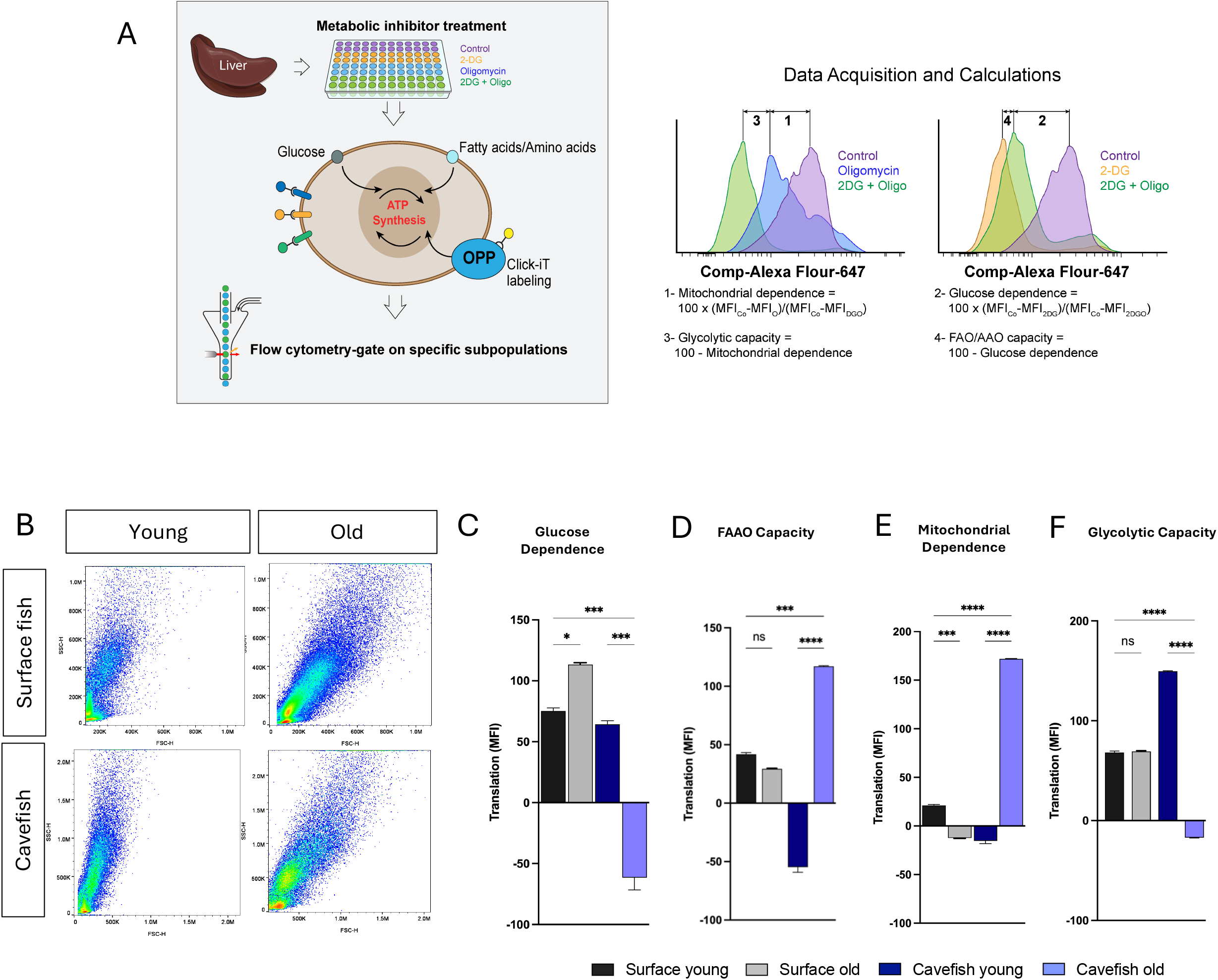
Metabolic reprogramming of energy source in old and young fish. A) SCENITH protocol adapted analyze dependence on diEerent ATP synthesis pathway in liver cells. (B) Cytometry data on liver cell populations measuring viable liver cells in young and old cavefish. (C-F) Bioenergetic analysis showing that pathway dependence varies between young and old fish in both surface and cavefish. n = 3, for surface and cavefish per timepoint. Data are represented as mean + SD, *P < 0.05, **P < 0.01, ***P < 0.001, ****P < 0.0001, ns = not significant.

SCENITH technology revealed that surface fish reliance on glucose significantly increased with age (**Figure 5C**) while FAAO decreased significantly in older fish (**Figure 5D**). Under normal conditions, glucose and fatty acid metabolism are synchronized to provide ATP through oxidative metabolism. However, during aging, metabolism is dramatically remodeled from oxidative process to anaerobic metabolism (Smith et al., 2018). This glucose related increase in cellular anaerobic metabolism with age, has been linked to increased oxidative stress and tissue damage (Chia et al., 2018; Saisho, 2014). Although mitochondrial dependence (**Figure 5E**) and glycolytic capacity (**Figure 5F**) did not change between young and old surface fish, the heightened glucose demand, coupled with elevated serum levels of glucose and lipid metabolism proteins, may correspond with increased anaerobic energy production. This shift could contribute to oxidative stress and the age-related effects observed in surface fish. ≠

In contrast, cavefish exhibited a reduced dependence on glucose and glycolytic capacity with age. Notably, mitochondrial activity and fatty acid oxidation (FAAO) were significantly elevated compared to younger fish. Since fats are exclusively oxidized to produce ATP (Voet & Voet, 2010), this enhanced oxidation and increased mitochondrial metabolism could imply increase energy production in aged cavefish, to support essential biological processes. However, given that reactive oxygen species (ROS) production is unavoidable, and continuous mitochondrial functioning inevitably leads to ROS generation (Bhatti et al., 2017; Murphy, 2009), the increased serum levels of anti-inflammatory and antioxidant adipokines in aged cavefish appears to counteract ROS production. Moreover, the increase in fatty acid oxidation could potentially fuel the pentose phosphate pathway (Carracedo et al., 2013; Lee et al., 2015), scavenging ROS and bolstering antioxidant defense mechanisms.

Together, these findings reinforce that the cavefish have finely modulated bioenergetics at the cellular level, which are crucial for maintaining optimal function and mitigating age-related stress and tissue damage. In contrast, surface fish lack these protective mechanisms, leading to more significant deterioration with age.

### Conclusion and limitation of study

Cavefish, despite their altered physiology, can live long, healthy lives, raising the question of whether they exhibit signs of aging. Our study reveals that aged surface fish show clear signs of aging, including spinal deformities, reduced body weight and impaired swimming ability, while cavefish do not display these indicators. We found that insulin signaling has limited contribution to the extended lifespan of cavefish. Notably, differences in energy source reprogramming and molecular signatures in circulating proteins possibly explained the more pronounced aging markers in surface fish. These findings highlight the resilience of cavefish to age-related changes despite their unusual physiology. While we have made progress in understanding cellular and molecular changes associated with aging, their interaction with metabolic regulators still needs more exploration. Indeed, this model offers a valuable opportunity to identify novel biomarkers and therapeutic strategies for promoting healthy aging and metabolic rejuvenation.

## Methods

### Ethics statement

All animal studies and methods were performed in accordance with approved animal protocols from the institutional Animal Care and Use Committees (qaaaaas) at the Stowers Institute for Medical Research. All methods described here for *Astyanax* experiments were approved on protocol 2024-175, while zebrafish were approved on protocol 2024-168. Housing conditions meet federal regulations and are accredited by AAALAC International.

### *Astyanax* husbandry

Surface fish, and Pachón cavefish morphs of *A. mexicanus* maintained under the same laboratory conditions were used in this study. They were kept in polycarbonate recirculating aquaculture racks (Pentair Aquatic Eco-Systems), with a 14:10-h light:dark photoperiod. Each rack system is equipped with mechanical, chemical, and biological filtration and UV disinfection. Adult fish were fed three times per day during non-breeding weeks and one time per day during breeding weeks on a diet of Mysis shrimp (Hikari Sales USA) and Gemma 800 (Skretting USA). Water quality parameters are monitored daily and were conducted similar to (Xiong et al., 2021).

### Zebrafish husbandry

Zebrafish housed in polycarbonate fish tanks were placed on racks (Pentair Aquatic Eco-Systems) with a 14/10 h light/dark photoperiod. Racks are supplied by two recirculating aquaculture systems with mechanical, chemical and biological filtration, and UV disinfection. Water quality parameters are maintained within safe limits (upper limit of total ammonia nitrogen range, 0.5 mg l−1; upper limit of nitrite range, 0.5 mg l−1; upper limit of nitrate range, 40 mg l−1; temperature set-point, 28.5 °C; pH, 7.60, specific conductance, 500 μS cm−1; dissolved oxygen, >85%). Water changes range from 20% to 30% daily (supplemented with Instant Ocean Sea Salt). Adult zebrafish are fed twice daily with one feed of hatched Artemia (first instar) (Brine Shrimp Direct) and one feed of Zeigler Adult Zebrafish Diet (Zeigler Bros). Embryos up to 5 days after fertilization were maintained at 28.5 °C in E2 embryo media.

### Morphological measurements

Body weight was obtained by placing animals dried on paper towels to remove excess water and weighing the animal on a tared scale. Other morphological measurements were obtained in ImageJ. Astyanax standard length was measured as a line drawn from the tip of the snout to the end of the central caudal fin ray (excluding the length of the caudal fin). Kyphosis angle was measured by taking the highest dorsal point of the outward curvature of the spine, a line drawn parallel to the standard-length line, and the angle between this line and the dorsal aspect of the caudal muscular was measured as the Kyphosis angle (Figure 1G)

### Zebrafish *insra* P211L genotyping

To genotype mutant animals, genomic DNA was extracted from zebrafish tail fin clips and used to amplify the *insra* locus, as previously described by (Riddle et al., 2018). Briefly, the following oligonucleotide primers were used, forward: GCACCCTTACACCCTTACATGA; and reverse: TACCGCTCAGCACTAATTTGGA; to target 700bp product size. PCR reactions containing 1× LA PCR Buffer II (Clontech), 2.5 mM MgCl^2^, 0.4 mM dNTP mix, 0.4 μM of each forward and reverse primer and 0.05 units of TaKaRa LA Taq DNA Polymerase (Clontech) were conducted in a 12.5-μl volume. The thermal cycling conditions were: initial denaturation at 94 °C for 2 minutes, followed by 35 cycles of 94 °C for 30 seconds, annealing temperature 52 °C for 30 seconds and 72 °C for 1 minute. A final elongation step was performed at 72 °C for 5 minutes. PCR products were diluted tenfold and sequenced directly on a 3730XL DNA Analyzer (Applied Biosystems) using the sequencing primer GGTGGAGTTGATGGTGGTATAG.

### Video Tracking and swim performance

Three surface fish and cavefish were placed individually in a 3-L water tank with continuous water circulation, modified to have internal acetal panels on front, bottom, and rear for improved video tracking. Fish were then recorded for 4 h beginning at the same time each day per experiment. The camera was positioned directly in front the tank. Prior to recording videos, animals were placed for 3 days to acclimatize to the arena. For each animal, digital video was captured at 30 frames per second and saved as mkv files. A fine-tuned Mask R-CNN model (He et al., 2017) was trained in Python using 250 frames randomly selected from the collections of 12 videos. The custom model was used to segment the fish in every frame to save the center-of-mass (x, y) and the bounding box of each fish. The (x, y) coordinates were used to calculate the velocity at each frame and the swimming distance. Calculations were performed in Python using numpy (Harris et al., 2020), pandas (Team, 2020), and scikit-image (van der Walt et al., 2014).

### Caudal fin regeneration assay on adult fish

For the caudal fin regeneration assay, both young and old cavefish and surface fish were anesthetized using 500 mg/L MS222. Approximately 2 mm of fin tissue was removed from the base of the caudal peduncle using a razor blade. Images of the cavefish fins were captured before amputation and at various days post-amputation (dpa) according to the experimental design. The fin tissue was preserved and processed as required for each experiment. Once the fish recovered from the procedure, they were returned to recirculating water for the remainder of the experiment. Each fish was individually housed, tracked and imaged to monitor regeneration progress over time. The percentage of fin regeneration was calculated by dividing the area of regrowth by the original fin area.

### Telomere length experiment and qPCR

Telomere length was measured from the caudal fin of young and old surface fish and cavefish by using real time quantitative PCR (Lin et al., 2019; Lindrose et al., 2021). A few millimeters from the ventral lobe of the caudal fin were collected and immediately stored in -70^°^C or stored in 80µl of lysis buffer (Promega). 50mg of fin samples were homogenized in Z742682 BeadBug™ 6 microtube homogenizer (Merck KGaA, Darmstadt, Germany) with ceramic beads. DNA was extracted following the Promega Maxwell® RSC Tissue DNA isolation Kit. We used the following primers to measure telomere length (Lunghi & Bilandzija, 2024). TEL – Forward (F) 5′-CGGTTTGTTTGGGTTTGGGT TTGGGTTTGGGTTTGGGTT-3′, Reverse (R) 5-GGCTTGCCTTACCCTTACCCTTACCCTTACCCTTACCCT-3′ and single copy gene used as reference (OCA2; forward (F) CAAGAACACTCTGGAGATGGAG, reverse (R) ACGCAGCTCGTCAAAGTT).

Primers were designed in the second exon and the amplicon was 109 bp in length. PCR was carried out using SYBR Green PCR Master Mix (Applied Biosystems). The expression levels of TEL were normalized to the levels of a single-copy gene oca2 (ENSAMXG00000012753) following recommendations of (Vasilishina et al., 2019) as allows us to estimate the relative average telomere length. All qRT-PCR assays were undertaken at the same time under identical conditions and performed in triplicate.

### Proteomics sample preparation

Serum samples collected from animals fasted overnight and immediately dissected following euthanization via submersion in ms222 (500mg/L), were frozen in liquid nitrogen, and stored at -80°C until further processing. For sample preparation, two-phase metabolite extraction was employed using methanol and chloroform. The non-extractable residue was collected and speed-vac to dryness. Pellets were reconstituted in buffer containing 90ul 8M urea, 100mM Tris-HCl pH 8.5, incubated with 5 mM TCEP (tris(2-carboxyethyl) phosphine) vortexed, sonicated and incubated in a thermomixer for 30 minutes at 37°C prior to enzymatic digestion. Digestion was done using 0.1 μg endoproteinase Lys-C or 6 hours at 37°C. The concentration of urea was reduced to 2 M using 100 mM Tris-HCl, pH 8.5. CaCl2 was added to 2 mM final concentration. Samples were incubated at 37°C overnight with 0.5 μg trypsin shaking and reactions were quenched with formic acid (5% final concentration).

### Mass spectrometry analysis and peptide detection

100ul digest was loaded onto a 3-phase column and eluted from the column using a 10-step Multidimensional protein identification technology (MudPIT) as previously described (Swanson et al., 2009). Mass spectrometry was performed using an Orbitrap Elite Hybrid mass spectrometer in positive ion mode. Resulting .raw files were converted to .ms2 files using the in-house software package RAWDistiller v. 1.0. The ProLuCID search algorithm was used to match spectra to a database containing 39383 *Astyanax mexicanus* protein sequences NCBI (NCBI 02/26/2021 release) and 426 common contaminant protein sequences. The result files from the ProLuCID search engine were processed with DTASelect (v 1.9) (Tabb et al., 2002) to assemble peptide level information into protein level information. Our in-house software, swallow and sandmartin (v 0.0.1), worked with DTASelect to select Peptide Spectrum Matches such as the FDRs at the peptide and protein levels were less than 1%.

Peptides and Proteins detected in these samples were compared using CONTRAST (Tabb et al., 2002). By combining all runs, we applied filtration criteria of fold change > 2.0 and P value < 0.05 (proteins had to be detected by at least 2 peptides) to identify significantly differentially expressed proteins (DEPs). Proteins that were subsets of others were removed using the parsimony option in DTASelect after merging all runs. Proteins that were identified by the same set of peptides (including at least one peptide unique to such protein group to distinguish between isoforms) were grouped together, and one accession number was arbitrarily considered as representative of each protein group. Protein abundances across all runs and samples were assessed by calculating the normalized spectral abundance factor (NSAF) using our in-house quantitative software (NSAF7, v 0.0.1) (Zhang et al., 2010). FDRs calculated by NSAF7 for these samples were less than 1% at all levels.

### ATP Generation Pathway Analysis

Investigating ATP generation was performed similarly to the SCENITH assay as described in (Arguello et al., 2020), except that the Click-iT™ Plus OPP Alexa Fluor™ 647 Protein Synthesis Assay Kit (ThermoFisher Scientific, Cat#: 10458) was used. Oligomycin (Fisher Scientific, Cat#: 41-105) and 2-deoxy-glucose (Fisher Scientific, Cat#: AC111980050) were bought separately. Briefly, isolated liver single cells were harvested and homogenized into 2mls of 1x PBS and filtered through a 70 µm mesh filter. Isolated cells were then separated into four groups of cells treated with drug inhibitors using the final concentration as follows: control (20 µM OPP (C)), oligomycin (20 µM OPP and 1 µM Oligomycin A (O)), 2-deoxy-glucose (20 µM OPP and 100 mM 2-deoxy-glucose (DG)), and combination of 2DG and O (2DGO) in addition to OPP. Cells were incubated thus for 30 minutes at 37°C. Subsequently, cells were washed, fixed with 3.7% PFA and permeabilized with 0.5% Triton X-100 OPP integration was then measured using the picolyl azide conjugated to Alexa Fluor 647 (AF647) according to the Click-iT™ Plus OPP Alexa Fluor™ 647 Protein Synthesis Assay Kit protocol. Cells were re-suspended in 1X PBS containing 0.1 µg/mL DAPI.

Finally, data acquisition was performed using the Cytek Aurora flow cytometer (Cytek) with high-throughput sampler. Unstained cell controls were used for autofluorescence extraction for each time point and metabolic inhibitor treatments (C, 2DG, O, DGO). Samples were unmixed using reference controls using the SpectroFlo Software v2.2.0.1. The unmixed FCS files were used for data processing and analysis using FlowJo v10.0.0. Cells were gated on FSC-A/SSC-A to gate out debris, then on FSC-A/FSC-H to identify single-cells and lastly on DAPI+ AF647+, and mean fluorescence intensity (MFI) of OPP AF647 in this population was then measured. Comparisons in OPP AF647 MFI was then performed by exporting data into GraphPad Prism. Calculations used to derive SCENITH parameters then performed according to the methods described by Arguello *et al*.

### Quantification and Statistical analysis

Data are presented as mean ± standard error of the mean (SEM) as marked unless specified differently. Multiple group comparison statistical significance was analyzed using an ordinary one-way or two-way ANOVA where appropriate, followed by Sidak’s or Dunnett’s correction. Zebrafish survival was plotted using Kaplan–Meier survival curves, statistical significance was determined via the log-rank test. P-values <0.05 were considered statistically significant represented as *p < 0.05, **p < 0.01, ***p < 0.001. The number of animals included in each experiment are indicated in the figure legend. Except for the proteomic analysis, all experiments were performed in at least 2 independent experiments. All statistics were executed using the GraphPad Prism software package (10.3.1).

## Acknowledgements

We are grateful to the Cavefish facility and Zebrafish facilities at the Stowers Institute for Medical Research for animal husbandry support. The authors would also like to thank Y. Hao for help in processing and acquisition of proteomics data, as well as Mark Miller for help with illustrations. Finally, we thank members of the Rohner lab, past and present for creating a motivating scientific environment and their stimulating discussions on the project. This work was supported by institutional funding from the Stowers Institute for Medical Research. Additionally, NR is supported by the National Institutes of Health (NIH) grant no. 1DP2AG071466-01 and R24OD030214.

## Author contributions

Conceptualization, A.E.C and N.R; methodology, A.E.C, J.E.J and N.R, investigation, A.E.C, A.K, J.E.J, S.S, P.M-S and C.W; analysis, A.E.C, A.K, J.E.J, S.S and C.W; validation, writing – original draft, A.E.C and N.R; writing – review and editing, A.E.C and N.R; supervision, A.E.C and N.R; resources, N.R; funding acquisition, N.R.

## Data depository

Original data underlying this manuscript may be accessed from the Stowers Original Data Repository at https://www.stowers.org/research/publications/libpb-2496 and/or are available from the corresponding author on reasonable request.

## REFERENCES

Anchelin, M., Murcia, L., Alcaraz-Perez, F., Garcia-Navarro, E. M., & Cayuela, M. L. (2011, Feb 9). Behaviour of telomere and telomerase during aging and regeneration in zebrafish. PLoS One, 6(2), e16955. 10.1371/journal.pone.0016955

Arguello, R. J., Combes, A. J., Char, R., Gigan, J. P., Baaziz, A. I., Bousiquot, E., Camosseto, V., Samad, B., Tsui, J., Yan, P., Boissonneau, S., Figarella-Branger, D., Gatti, E., Tabouret, E., Krummel, M. F., & Pierre, P. (2020, Dec 1). SCENITH: A Flow Cytometry-Based Method to Functionally Profile Energy Metabolism with Single-Cell Resolution. Cell Metab, 32(6), 1063–1075 e1067. 10.1016/j.cmet.2020.11.007

Barzilai, N., Huffman, D. M., Muzumdar, R. H., & Bartke, A. (2012, Jun). The critical role of metabolic pathways in aging. Diabetes, 61(6), 1315–1322. 10.2337/db11-1300

Bhatti, J. S., Bhatti, G. K., & Reddy, P. H. (2017, May). Mitochondrial dysfunction and oxidative stress in metabolic disorders - A step towards mitochondria based therapeutic strategies. Biochim Biophys Acta Mol Basis Dis, 1863(5), 1066–1077. 10.1016/j.bbadis.2016.11.010

Blackburn, E. H., Greider, C. W., & Szostak, J. W. (2006, Oct). Telomeres and telomerase: the path from maize, Tetrahymena and yeast to human cancer and aging. Nat Med, 12(10), 1133–1138. 10.1038/nm1006-1133

Boonekamp, J. J., Simons, M. J., Hemerik, L., & Verhulst, S. (2013, Apr). Telomere length behaves as biomarker of somatic redundancy rather than biological age. Aging Cell, 12(2), 330–332. 10.1111/acel.12050

Bratic, I., & Trifunovic, A. (2010). Mitochondrial energy metabolism and ageing. Biochimica et Biophysica Acta (BBA)-Bioenergetics, 1797(6-7), 961–967.

Camilli, C., Hoeh, A. E., De Rossi, G., Moss, S. E., & Greenwood, J. (2022, Jan 21). LRG1: an emerging player in disease pathogenesis. J Biomed Sci, 29(1), 6. 10.1186/s12929-022-00790-6

Carracedo, A., Cantley, L. C., & Pandolfi, P. P. (2013, Apr). Cancer metabolism: fatty acid oxidation in the limelight. Nat Rev Cancer, 13(4), 227–232. 10.1038/nrc3483

Chia, C. W., Egan, J. M., & Ferrucci, L. (2018, Sep 14). Age-Related Changes in Glucose Metabolism, Hyperglycemia, and Cardiovascular Risk. Circ Res, 123(7), 886–904. 10.1161/CIRCRESAHA.118.312806

Choi, C. H. J., Barr, W., Zaman, S., Model, C., Park, A., Koenen, M., Lin, Z., Szwed, S. K., Marchildon, F., Crane, A., Carroll, T. S., Molina, H., & Cohen, P. (2022, Nov 8). LRG1 is an adipokine that promotes insulin sensitivity and suppresses inflammation. Elife, 11, e81559. 10.7554/eLife.81559

Cobham, A. E., & Rohner, N. (2024, May). Unraveling stress resilience: Insights from adaptations to extreme environments by Astyanax mexicanus cavefish. J Exp Zool B Mol Dev Evol, 342(3), 178–188. 10.1002/jez.b.23238

Davidson, R. T., Arias, E. B., & Cartee, G. D. (2002, Feb). Calorie restriction increases muscle insulin action but not IRS-1-, IRS-2-, or phosphotyrosine-PI 3-kinase. Am J Physiol Endocrinol Metab, 282(2), E270–276. 10.1152/ajpendo.00232.2001

Ebert, T. A. (1982). Longevity, life history, and relative body wall size in sea urchins. Ecological Monographs, 52(4), 353–394.

Elmore, L. W., Norris, M. W., Sircar, S., Bright, A. T., McChesney, P. A., Winn, R. N., & Holt, S. E. (2008, Aug). Upregulation of telomerase function during tissue regeneration. Exp Biol Med (Maywood), 233(8), 958–967. 10.3181/0712-RM-345

Fried, L. P., Ferrucci, L., Darer, J., Williamson, J. D., & Anderson, G. (2004, Mar). Untangling the concepts of disability, frailty, and comorbidity: implications for improved targeting and care. J Gerontol A Biol Sci Med Sci, 59(3), 255–263. 10.1093/gerona/59.3.m255

Gerhard, G. S., Kauffman, E. J., Wang, X., Stewart, R., Moore, J. L., Kasales, C. J., Demidenko, E., & Cheng, K. C. (2002, Aug-Sep). Life spans and senescent phenotypes in two strains of Zebrafish (Danio rerio). Exp Gerontol, 37(8-9), 1055–1068. 10.1016/s0531-5565(02)00088-8

Gorbunova, V., Bozzella, M. J., & Seluanov, A. (2008, Sep). Rodents for comparative aging studies: from mice to beavers. Age, 30(2-3), 111–119. 10.1007/s11357-008-9053-4

Gresl, T. A., Colman, R. J., Havighurst, T. C., Byerley, L. O., Allison, D. B., Schoeller, D. A., & Kemnitz, J. W. (2003, Dec). Insulin sensitivity and glucose effectiveness from three minimal models: effects of energy restriction and body fat in adult male rhesus monkeys. Am J Physiol Regul Integr Comp Physiol, 285(6), R1340–1354. 10.1152/ajpregu.00651.2002

Harman, D. (1972, Apr). The biologic clock: the mitochondria? J Am Geriatr Soc, 20(4), 145–147. 10.1111/j.1532-5415.1972.tb00787.x

Harris, C. R., Millman, K. J., van der Walt, S. J., Gommers, R., Virtanen, P., Cournapeau, D., Wieser, E., Taylor, J., Berg, S., Smith, N. J., Kern, R., Picus, M., Hoyer, S., van Kerkwijk, M. H., Brett, M., Haldane, A., Del Río, J. F., Wiebe, M., Peterson, P., Gérard-Marchant, P., Sheppard, K., Reddy, T., Weckesser, W., Abbasi, H., Gohlke, C., & Oliphant, T. E. (2020, Sep). Array programming with NumPy. Nature, 585(7825), 357–362. 10.1038/s41586-020-2649-2

He, K., Gkioxari, G., Dollár, P., & Girshick, R. (2017). Mask r-cnn. Proceedings of the IEEE international conference on computer vision,

Healey, M. (1991). Life history of chinook salmon (Oncorhynchus tshawytscha). Pacific salmon life histories, 311–394.

Kalant, N., Stewart, J., & Kaplan, R. (1988, Dec). Effect of diet restriction on glucose metabolism and insulin responsiveness in aging rats. Mech Ageing Dev, 46(1-3), 89–104. 10.1016/0047-6374(88)90117-0

Kenyon, C. (2005, Feb 25). The plasticity of aging: insights from long-lived mutants. Cell, 120(4), 449–460. 10.1016/j.cell.2005.02.002

Kenyon, C., Chang, J., Gensch, E., Rudner, A., & Tabtiang, R. (1993). A C. elegans mutant that lives twice as long as wild type. Nature, 366(6454), 461–464. 10.1038/366461a0

Lee, E. A., Angka, L., Rota, S. G., Hanlon, T., Mitchell, A., Hurren, R., Wang, X. M., Gronda, M., Boyaci, E., Bojko, B., Minden, M., Sriskanthadevan, S., Datti, A., Wrana, J. L., Edginton, A., Pawliszyn, J., Joseph, J. W., Quadrilatero, J., Schimmer, A. D., & Spagnuolo, P. A. (2015, Jun 15). Targeting Mitochondria with Avocatin B Induces Selective Leukemia Cell Death. Cancer Res, 75(12), 2478–2488. 10.1158/0008-5472.CAN-14-2676

Lopez-Otin, C., Blasco, M. A., Partridge, L., Serrano, M., & Kroemer, G. (2013, Jun 6). The hallmarks of aging. Cell, 153(6), 1194–1217. 10.1016/j.cell.2013.05.039

Lopez-Otin, C., Blasco, M. A., Partridge, L., Serrano, M., & Kroemer, G. (2023, Jan 19). Hallmarks of aging: An expanding universe. Cell, 186(2), 243–278. 10.1016/j.cell.2022.11.001

Lowe, D. A., Degens, H., Chen, K. D., & Alway, S. E. (2000, Mar). Glyceraldehyde-3-phosphate dehydrogenase varies with age in glycolytic muscles of rats. J Gerontol A Biol Sci Med Sci, 55(3), B160–164. 10.1093/gerona/55.3.b160

Lund, T. C., Glass, T. J., Tolar, J., & Blazar, B. R. (2009, Nov 6). Expression of telomerase and telomere length are unaffected by either age or limb regeneration in Danio rerio. PLoS One, 4(11), e7688. 10.1371/journal.pone.0007688

Lunghi, E., & Bilandzija, H. (2024). Telomere length and dynamics in Astyanax mexicanus cave and surface morphs. PeerJ, 12, e16957. 10.7717/peerj.16957

McCarter, R., Mejia, W., Ikeno, Y., Monnier, V., Kewitt, K., Gibbs, M., McMahan, A., & Strong, R. (2007, Oct). Plasma glucose and the action of calorie restriction on aging. J Gerontol A Biol Sci Med Sci, 62(10), 1059–1070. 10.1093/gerona/62.10.1059

Medley, J. K., Persons, J., Biswas, T., Olsen, L., Peuss, R., Krishnan, J., Xiong, S., & Rohner, N. (2022, Jun 15). The metabolome of Mexican cavefish shows a convergent signature highlighting sugar, antioxidant, and Ageing-Related metabolites. Elife, 11. 10.7554/eLife.74539

Murphy, M. P. (2009). How mitochondria produce reactive oxygen species. Biochemical journal, 417(1), 1–13.

Nicholls, C., Pinto, A. R., Li, H., Li, L., Wang, L., Simpson, R., & Liu, J. P. (2012, Aug 14). Glyceraldehyde-3-phosphate dehydrogenase (GAPDH) induces cancer cell senescence by interacting with telomerase RNA component. Proc Natl Acad Sci U S A, 109(33), 13308–13313. 10.1073/pnas.1206672109

Olsen, L., Levy, M., Medley, J. K., Hassan, H., Miller, B., Alexander, R., Wilcock, E., Yi, K., Florens, L., Weaver, K., McKinney, S. A., Peuss, R., Persons, J., Kenzior, A., Maldonado, E., Delventhal, K., Gluesenkamp, A., Mager, E., Coughlin, D., & Rohner, N. (2023, Jan 31). Metabolic reprogramming underlies cavefish muscular endurance despite loss of muscle mass and contractility. Proc Natl Acad Sci U S A, 120(5), e2204427120. 10.1073/pnas.2204427120

Olsen, L., Thum, E., & Rohner, N. (2021, May 17). Lipid metabolism in adaptation to extreme nutritional challenges. Dev Cell, 56(10), 1417–1429. 10.1016/j.devcel.2021.02.024

Park, H. N., Song, M. J., Choi, Y. E., Lee, D. H., Chung, J. H., & Lee, S. T. (2023, Aug 4). LRG1 Promotes ECM Integrity by Activating the TGF-beta Signaling Pathway in Fibroblasts. Int J Mol Sci, 24(15), 12445. 10.3390/ijms241512445

Patch, A., Paz, A., Holt, K. J., Duboue, E. R., Keene, A. C., Kowalko, J. E., & Fily, Y. (2022). Kinematic analysis of social interactions deconstructs the evolved loss of schooling behavior in cavefish. PLoS One, 17(4), e0265894. 10.1371/journal.pone.0265894

Paz, A., McDole, B., Kowalko, J. E., Duboue, E. R., & Keene, A. C. (2020, Nov). Evolution of the acoustic startle response of Mexican cavefish. J Exp Zool B Mol Dev Evol, 334(7-8), 474–485. 10.1002/jez.b.22988

Riddle, M. R., Aspiras, A., Damen, F., McGaugh, S., Tabin, J. A., & Tabin, C. J. (2021, May 21). Genetic mapping of metabolic traits in the blind Mexican cavefish reveals sex-dependent quantitative trait loci associated with cave adaptation. BMC Ecol Evol, 21(1), 94. 10.1186/s12862-021-01823-8

Riddle, M. R., Aspiras, A. C., Gaudenz, K., Peuss, R., Sung, J. Y., Martineau, B., Peavey, M., Box, A. C., Tabin, J. A., McGaugh, S., Borowsky, R., Tabin, C. J., & Rohner, N. (2018, Mar 29). Insulin resistance in cavefish as an adaptation to a nutrient-limited environment. Nature, 555(7698), 647–651. 10.1038/nature26136

Saisho, Y. (2014). Glycemic Variability and Oxidative Stress: A Link between Diabetes and Cardiovascular Disease? International Journal of Molecular Sciences, 15(10), 18381–18406. https://www.mdpi.com/1422-0067/15/10/18381

Shilovsky, G. A., Putyatina, T. S., & Markov, A. V. (2022, Dec). Evolution of Longevity as a Species-Specific Trait in Mammals. Biochemistry (Mosc), 87(12), 1579–1599. 10.1134/S0006297922120148

Smith, R. L., Soeters, M. R., Wust, R. C. I., & Houtkooper, R. H. (2018, Aug 1). Metabolic Flexibility as an Adaptation to Energy Resources and Requirements in Health and Disease. Endocr Rev, 39(4), 489–517. 10.1210/er.2017-00211

Swanson, S. K., Florens, L., & Washburn, M. P. (2009). Generation and analysis of multidimensional protein identification technology datasets. Methods Mol Biol, 492, 1–20. 10.1007/978-1-59745-493-3_1

Tabb, D. L., McDonald, W. H., & Yates, J. R., 3rd. (2002, Jan-Feb). DTASelect and Contrast: tools for assembling and comparing protein identifications from shotgun proteomics. J Proteome Res, 1(1), 21–26. 10.1021/pr015504q

Tan, D., Patton, P., & Coombs, S. (2011, Jul). Do blind cavefish have behavioral specializations for active flow-sensing? J Comp Physiol A Neuroethol Sens Neural Behav Physiol, 197(7), 743–754. 10.1007/s00359-011-0637-6

Tatar, M., Bartke, A., & Antebi, A. (2003, Feb 28). The endocrine regulation of aging by insulin-like signals. Science, 299(5611), 1346–1351. 10.1126/science.1081447

Team, T. P. D. (2020). pandas-dev/pandas: Pandas. Zenodo, February.

Thomas, D. R. (2007, Aug). Loss of skeletal muscle mass in aging: examining the relationship of starvation, sarcopenia and cachexia. Clin Nutr, 26(4), 389–399. 10.1016/j.clnu.2007.03.008

Tian, Y., Tian, Y., Yuan, Z., Zeng, Y., Wang, S., Fan, X., Yang, D., & Yang, M. (2022, Mar 25). Iron Metabolism in Aging and Age-Related Diseases. Int J Mol Sci, 23(7), 3612. 10.3390/ijms23073612

van der Walt, S., Schönberger, J. L., Nunez-Iglesias, J., Boulogne, F., Warner, J. D., Yager, N., Gouillart, E., & Yu, T. (2014). scikit-image: image processing in Python. PeerJ, 2, e453. 10.7717/peerj.453

Vasilishina, A., Kropotov, A., Spivak, I., & Bernadotte, A. (2019). Relative Human Telomere Length Quantification by Real-Time PCR. Methods Mol Biol, 1896, 39–44. 10.1007/978-1-4939-8931-7_5

Vaupel, J. W., Manton, K. G., & Stallard, E. (1979, Aug). The impact of heterogeneity in individual frailty on the dynamics of mortality. Demography, 16(3), 439–454. https://www.ncbi.nlm.nih.gov/pubmed/510638

Voet, D., & Voet, J. G. (2010). Biochemistry. John Wiley & Sons.

Xiong, S., Wang, W., Kenzior, A., Olsen, L., Krishnan, J., Persons, J., Medley, K., Peuß, R., Wang, Y., Chen, S., Zhang, N., Thomas, N., Miles, J. M., Sánchez Alvarado, A., & Rohner, N. (2021). Enhanced lipogenesis through Pparγ helps cavefish adapt to food scarcity. bioRxiv. 10.1101/2021.04.27.441667

Zhang, Y., Wen, Z., Washburn, M. P., & Florens, L. (2010, Mar 15). Refinements to label free proteome quantitation: how to deal with peptides shared by multiple proteins. Anal Chem, 82(6), 2272–2281. 10.1021/ac9023999

